# ASD Diagnosis in Adults: Phenotype and Genotype Findings from a Clinically-derived Cohort

**DOI:** 10.1101/420778

**Authors:** Underwood Jack F G, Kendall Kimberley M, Berrett Jennifer, Anney Richard, Van den Bree Marianne B.M., Hall Jeremy

## Abstract

**Background:** The last decade has seen the development of services for adults presenting with symptoms of autism spectrum disorder (ASD) in the UK. Compared to children, little is known about the phenotypic and genetic characteristics of these patients.

**Aims:** This e-cohort study aimed to examine the phenotypic and genetic characteristics of a clinically-presenting sample of adults diagnosed with ASD by specialist services.

**Methods:** Individuals diagnosed with ASD as adults were recruited by the National Centre for Mental Health and completed self-report questionnaires, interviews and provided DNA. 105 eligible individuals were matched to 76 healthy controls. We investigated the demographics, social history, comorbid psychiatric and physical disorders. Samples were genotyped, copy number variants (CNVs) were called and polygenic risk scores calculated.

**Results:** 89.5% of individuals with ASD had at least one comorbid psychiatric diagnosis with comorbid depression (62.9%) and anxiety (55.2%) the most common. The ASD group experienced more neurological comorbidities than healthy controls, particularly migraine headache. They were less likely to have married or be in work and had more alcohol-related problems. There was a significantly higher load of autism common genetic variants in the adult ASD group compared to controls, but there was no difference in the rate of rare CNVs.

**Conclusions:** This study provides important information about psychiatric comorbidity in adult ASD which may be used to inform clinical practice and patient counselling. It also suggests that the polygenic load of common ASD-associated variants may be important in conferring risk within non-intellectually disabled population of adults with ASD.

## Introduction

Autism Spectrum Disorder (ASD) is a group of neurodevelopmental disorders characterised by persistent difficulties in social interaction and communication, as well as restricted interests, stereotypic behaviours and resistance to change^1^. Epidemiological studies report a prevalence of ASD in the general population of around 1%, with a male to female ratio of approximately 3:1 ^2–5^. The majority of studies of ASD have been carried out in paediatric populations and have included individuals with intellectual disability (ID). The limited studies in individuals diagnosed with ASD as adults have shown that adults with ASD experience significant disadvantage in employment, social relationships and quality of life ^6,7^. In these studies, ASD has also been associated with increased lifetime psychiatric comorbidity ^8–10^. There has been very little research reporting lifetime outcomes for individuals diagnosed with ASD as adults. This information has the potential to inform specialist service development and tailor clinical care. Advancing knowledge in this area is important as following the Autism Act in 2009 and recommendation in the NICE Guidelines of 2012 many adult ASD diagnostic services have been set up across the UK, although provision remains sporadic ^11–13^. Furthermore, while some adult services now offer genetic testing for individuals diagnosed with ASD, little is known about the genetic characteristics of adults presenting with ASD. Studies of predominantly paediatric ASD populations have shown a substantial contribution of both rare copy number variants (CNVs) and polygenic burden of common variants to risk for ASD. Rare CNVs are reported to occur in 10-15% of childhood ASD cases, a finding which has encouraged formal genetic testing in childhood ASD ^14,15^. However, it is not known whether similar rates are seen in individuals presenting to adult diagnostic services. In this study we examine the demographic, social, psychiatric and physical health characteristics of a cohort of individuals presenting with ASD in adulthood compared to a healthy control population from the same source databank. We also report the rates of neurodevelopmental CNVs and polygenic burden of common variants associated with ASD in this sample.

## Methods

### Sample

Data was obtained from the National Centre for Mental Health (NCMH), a Welsh Government-funded Research Centre that investigates neurodevelopmental, adult, and neurodegenerative psychiatric disorders across the lifespan ^16^. Participants were recruited using a variety of systematic approaches in primary, secondary and tertiary health care services, including (a) the identification of potential participants by clinical care teams, (b) screening of clinical notes, and (c) the use of disease registers. The majority of participants in our sample were recruited via specialist diagnostic services. Non-systematic recruitment approaches included advertising in local media, placing posters and leaflets in NHS waiting areas, liaising with voluntary organisations and contacting individuals enrolled in previous studies within the Institute of Psychological Medicine and Clinical Neurosciences, Cardiff. To allow comparisons, the cohort includes control participants who self-report no experiences of any mental health disorder. All adult participants included in this study provided written informed consent for recruitment into NCMH. Trained research assistants administered a brief standardized interview assessment to consenting participants to ascertain details related to the participant’s personal and family history of mental health experiences including any comorbidity, past and current medication use, socio-demographic information including employment and education, physical health diagnoses and any substance misuse. A sample of venous blood or saliva was taken for genetic and other analyses. Participants were given a pack of standardised self-report questionnaires to complete and return to the research team by post after the initial assessment. Confirmation and further information regarding the primary diagnosis of ASD was obtained from clinical records where appropriate consent had been obtained to do so.

To date 10,870 individuals have been recruited to NCMH ^16^. At the point of initial search in June 2016 the database included ∼6,600 participants, of whom 172 individuals held a primary self-report diagnosis of Aspergers, Autism Spectrum Disorder or Autism and no self-report comorbid ID, and were potentially eligible for inclusion. On casenote review, 67 of the 172 participants were excluded, predominantly due to loss to follow-up (n=37, full details in Supplementary Material). The remaining 105 individuals were all confirmed to have an ASD diagnosis consistent with ICD-10 criteria by case-note review, with no evidence of recorded intellectual disability, and with a first diagnosis of ASD made by secondary care clinicians assessing in a diagnostic role when the participant was over the age of 18 ^1^. Seventy-six controls matched on age (within 5 years and over 18), ethnicity and sex, were obtained from the NCMH database. Controls participants were selected from individuals in the NCMH Database with no current or previous self-reported difficulties with mental health and no psychotropic medication usage. A favourable ethical opinion was received from the Wales Research Ethics Committee 2 on 25^th^ November 2016 for NCMH, and through internal NCMH applications for this study.

## Measures

### Demographic Information

Demographic data was collected by interview and questionnaire. Biological offspring was recorded in binary yes/ no variable and with free text number and biological age of children, recorded in 101 of the 105 ASD participants and all controls. Lifetime marriage and co-habitation was recorded as a binary yes/ no variable, recorded in 97 of the participants and all 76 controls. Current profession responses were multiple-choice skill level (see supplementary material for categorisation), reduced to ‘currently in work’/ ‘currently not in work’ for our analysis and recorded for 97 ASD participants and 73 controls.

### Comorbidities

Physical health comorbidities were established with a multiple-choice list of 22 common physical diagnoses (Supplementary Material), reduced to clinical system categories for comparison. Lifetime mental health comorbidities were established with a multiple-choice list of 37 common psychiatric diagnoses as reported in the supplementary material. By definition the control group had no psychiatric comorbidity for comparative analysis, thus preventing comparative analysis of mental health comorbid rates. Fifty-nine of the ASD participants and 53 of the control participants completed the Beck’s Depression Inventory (BDI) ^17^ providing information on current mood. Information on lifetime psychotropic medication usage was provided by 93-95 of the ASD participants.

### Substance Use

Individuals reported on problems encountered in their lifetime through substance use in financial, medical, relationship and occupational domains. For analysis purposes these were reduced to binary yes/no categories. Fifty-eight ASD participants and 60 of the 76 control participants responded to alcohol problem questions. Regular smoking was reported as a binary yes/no variable, as was regular cannabinoid use, with complete data available for 79 ASD participants and all controls. Regular cannabinoid use was reported as a binary yes/no variable and completed by 31 ASD participants and 25 control participants Usage of other street drugs was recorded through initial binary yes/no variable, and if positive, through a multiple-choice question incorporating common UK street drugs. Eighty ASD participants and all controls responded to the initial question, with 13 ASD participants and 13 controls giving further answers.

## Phenotype Statistical Analysis

Statistical analysis was performed utilising IBM SPSS Statistics 23 for Windows ^18^. Prevalence of comorbid psychiatric disorders was analysed through descriptive statistics for mean, standard deviation, variance and range. No comparative statistics of comorbid psychiatric diagnosis and associated medication usage was possible as by definition our control population was unaffected. Prevalence of sociodemographic, physical health comorbidity and family history was initially graphed. For normally distributed dependent variables analysis was by a chi-square test followed by a binomial logistic regression with ASD diagnosis (yes/no), age and sex entered as co-variants. Where dependent variables were non-normal, a non-parametric Mann-Whitney U test was used with Fisher’s exact test for expected cell counts <5.

## Genotyping

DNA was obtained from blood and saliva samples, and genotyping was carried out at the MRC Centre for Neuropsychiatric Genetics and Genomics, Hadyn Ellis Laboratory, Cardiff University, Wales. Samples were available from 90 individuals with ASD and 60 control participants and were genotyped on two versions of the Illumina Psych Chip - InfiniumPsycharray-24v1-2 (34 individuals with ASD, 59 controls) and InfiniumPsycharray with custom content (IPCMN Psych Chip – 56 individuals with ASD, 1 control). Analyses were based on the 90% common content between the arrays.

## CNV Calling

CNVs were called using PennCNV run through a custom Galaxy pipeline ^19,20^. Individual samples were excluded if they had 30 or more CNVs, had a waviness factor (WF) >0.03 or <-0.03 or a call rate <96%. Individual CNVs were excluded if their LRR Standard Deviation was >0.2. CNVs constituting less than 50kb or >10 single nucleotide polymorphisms (SNPs) were removed utilising a UNIX based script prior to annotation (Supplementary Material). Three hundred and seventy-three samples remained after QC. We annotated the CNVs called with a list of 53 CNVs associated with neurodevelopmental disorders ^21–23^ (Supplementary Material). The breakpoints of the initial call of CNVs were inspected to confirm they met the CNV calling criteria. We required a CNV to cover more than half the critical interval and to include key genes in the region (if known), or in the case of single gene CNVs we required deletions to intersect at least one exon and duplications to cover the whole gene ^23^.

## Polygenic Risk Scores

Raw genotype data underwent marker and individual quality control using the -genotypeqc-package. Individuals were excluded based on heterogeneity (4x deviation from sample standard deviation), excessive relatedness and missingness (>2%). Markers were excluded based on minor allele count (5), SNP missingness (>2%), Hardy-Weinberg equilibrium (p≤10-10) and deviation from reference MAF (>10%). Ancestry informative principle components were generated from linkage independent ancestry informative markers using the - bim2eigenvec-package. Training genome-wide association studies (GWAS) for Polygenic Risk Scores (PRS) were cleaned using the -summaryqc-package and processed using - summaryqc2prePRS-. All markers were mapped to hg19 and nomenclature standardised to the 1000 genomes reference panel based on chromosome location and UIPAC genotype code via the -snp2refid-package. PRS scores for ASD ^24^, ADHD ^25^, BIP ^26^, MDD ^27^, Alzheimer’s disease ^28^ and Schizophrenia ^29^ were generated for each individual using plink via the - profilescore_beta-. PRS were calculated for linkage independent markers (r^2 < 0.2) at association p-value thresholds of P< 0.5, P< 0.1, P< 0.05. P< 0.01, P< 1e-3, P< 1e-4, P< 1e-5, P< 1e-6, P< 1e-7, P< 1e-8. SNPs included in each model, q-score and score files are available on request. All packages used in this analysis are self-authored and implemented in Stata 13.0; packages and code available at https://github.com/ricanney/stata. The differences in the variance explained by each score for caseness, defined as the presence or absence of ASD (0 vs 1), was calculated using a model including the covariates allele count, sex and 10 ancestry derived principle components.

## Results

By definition control participants did not have psychiatric morbidity and were not using any psychotropic medication.

### Psychiatric Comorbidity

Comorbid psychiatric diagnosis was reported by 89.5% (n = 94) of individuals with ASD (Table 1). The most common comorbid diagnoses were depression (62.9%, n = 66) and anxiety (55.2%, n = 58). 44.8% of individuals with ASD reported dual depression and anxiety diagnoses. The Mean BDI score for the ASD participants was 20.356, which is on the border of mild (14-19) and moderate (20-28) depression severity. The mean for the control participants was 5.226, significantly different to the ASD participants (Z-score of −7.205, p < 0.001), and within the minimal range (0-13). The significant association was seen across all BDI subscale scores except Question 19 (weight change).

**Table 1.** Psychiatric comorbidity as defined by ICD-10 Reported psychiatric comorbidity in ASD participants grouped by WHO ICD-10 classification. *n* - number of individuals reporting the diagnosis; % - percentage of total who responded.

|  |  | ASD Participants (n=105) |
| --- | --- | --- |
|  | All comorbid psychiatric diagnosis | 94 (89.5%) |
| **F30-39**<br>Mood (affective) disorder | Any | 69 (65.7%) |
|  | Depression | 66 (62.9%) |
|  | Bipolar Disorder | 9 (8.6%) |
|  | Hypomania | 1 (1.0%) |
| **F40-48**<br>Neurotic, stress-related and somatoform | Any | 63 (60.0%) |
|  | Anxiety | 58 (55.2%) |
|  | OCD | 18 (17.1%) |
|  | Phobias | 9 (8.6%) |
|  | Agoraphobia | 8 (7.6%) |
|  | Panic Disorder | 8 (7.6%) |
|  | PTSD | 6 (5.7%) |
| **F80-89**<br>Psychological development | Any | 30 (28.6%) |
|  | Dyslexia | 23 (21.9%) |
|  | Dyspraxia | 19 (18.1%) |
| **F90-98**<br>Disorders with onset occurring in childhood or adolescence | Any | 22 (21.0%) |
|  | ADHD | 20 (19.0%) |
|  | Oppositional Defiant Disorder | 1 (0.6%) |
|  | Tic Disorders | 2 (1.9%) |
| **F20-29**<br>Schizophrenia, schizotypal and delusional disorders | Any | 13 (12.4%) |
|  | Psychosis | 12 (11.4%) |
|  | Schizophrenia | 4 (3.8%) |
|  | Schizoaffective Disorder | 3 (2.9%) |
| **F10-19**<br>Disorders due to substance use | Any | 13 (12.4%) |
|  | Alcohol Abuse | 10 (9.5%) |
|  | Other Substance Misuse/ Abuse | 9 (8.6%) |
| **F50-59**<br>Syndromes associated physiological disturbance | Any | 8 (7.6%) |
|  | Anorexia | 4 (3.8%) |
|  | Postnatal Depression | 3 (2.9%) |
|  | Bulimia | 2 (1.9%) |
| **F60-69**<br>Disorders of adult personality | Any | 4 (3.8%) |
|  | Borderline Personality Disorder | 3 (2.9%) |
|  | Other Personality Disorder | 1 (1.0%) |
| **F00-09**<br>Organic disorders | Pick's Disease (Frontotemporal Dementia) | 1 (1.0%) |

### Psychotropic Medication Usage

The most widely prescribed medications amongst ASD participants were antidepressants and anxiolytics and over a quarter reported taking antipsychotics (Table 2).

**Table 2.** Psychotropic medication usage amongst participants with ASD Reported psychotropic medication usage in participants with ASD based on NCMH questionnaire response fields. N - number of individuals reporting usage of medication of this type; % - percentage of total who responded.

|  | Reported lifetime use (n) |
| --- | --- |
| Stimulants | 6 (6.3%) |
| Antidepressants | 74 (78.7%) |
| Anxiolytics | 29 (31.2%) |
| Hypnotics | 17 (18.1%) |
| Mood Stabilisers | 6 (6.5%) |
| Antipsychotics | 25 (26.3%) |

### Sociodemographic phenotype

Data comparing sociodemographic phenotypes, physical health and family history for participants with ASD and controls is presented in Table 3. The average age of the ASD participants was 37.8 years (SD=12.3) and for controls it was 40.7 years (SD=14.1)(p=.89) due to matching. There were 80 males/ 25 females among the ASD participants (76.2% male) and 55 males/ 21 females in the control participants (72.4% male)(p=.56). Adults with ASD were significantly less likely to be currently working (OR=.174, p=<.001) or to be married or cohabiting (OR=.29, p=.002), to be currently off work due to sickness or disablement (OR=69.305, p=<.001) and to have alcohol-related problem (OR 6.24, p=.001). No differences were found in rates of having one or more biological children.

**Table 3.**
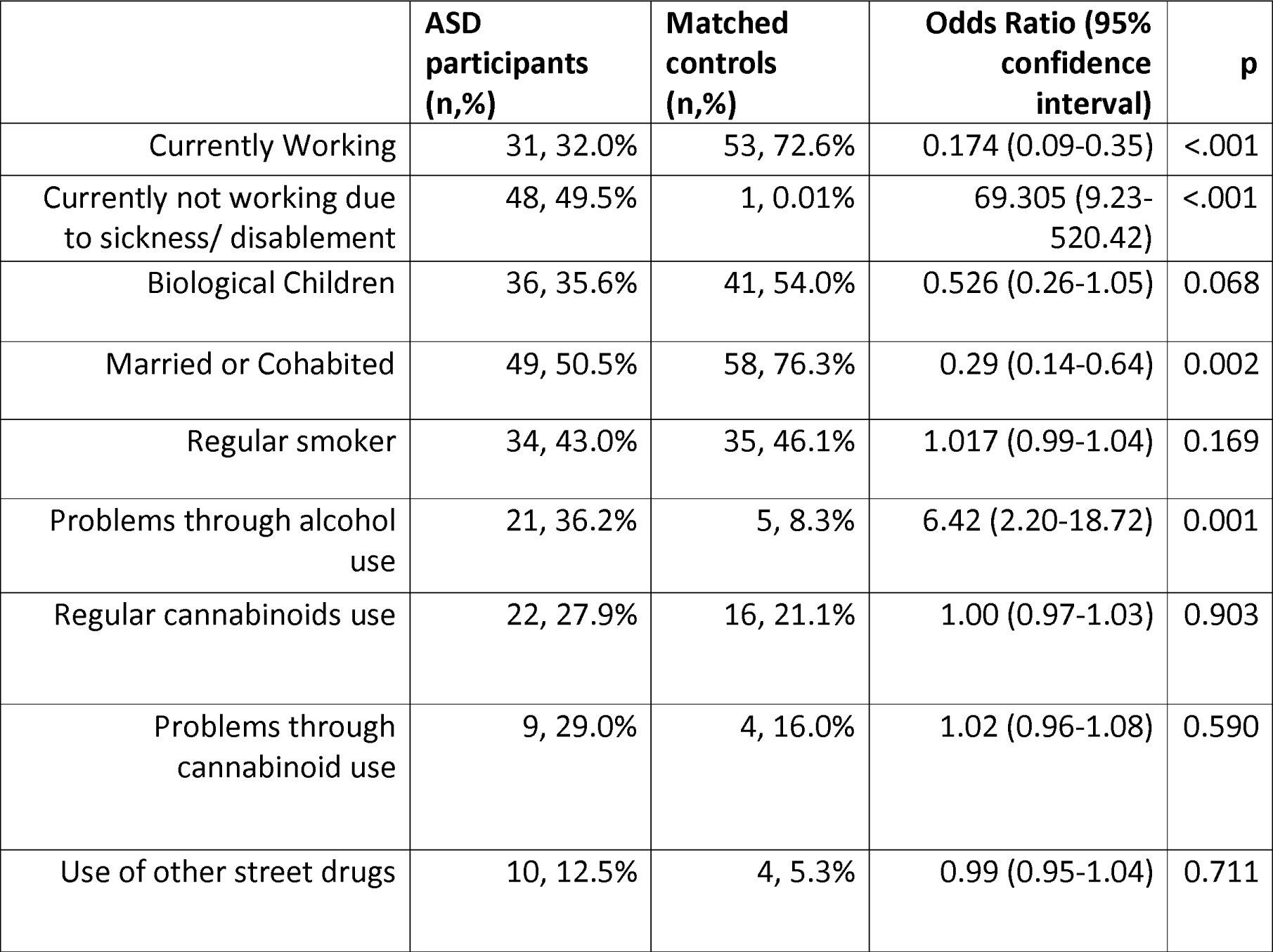
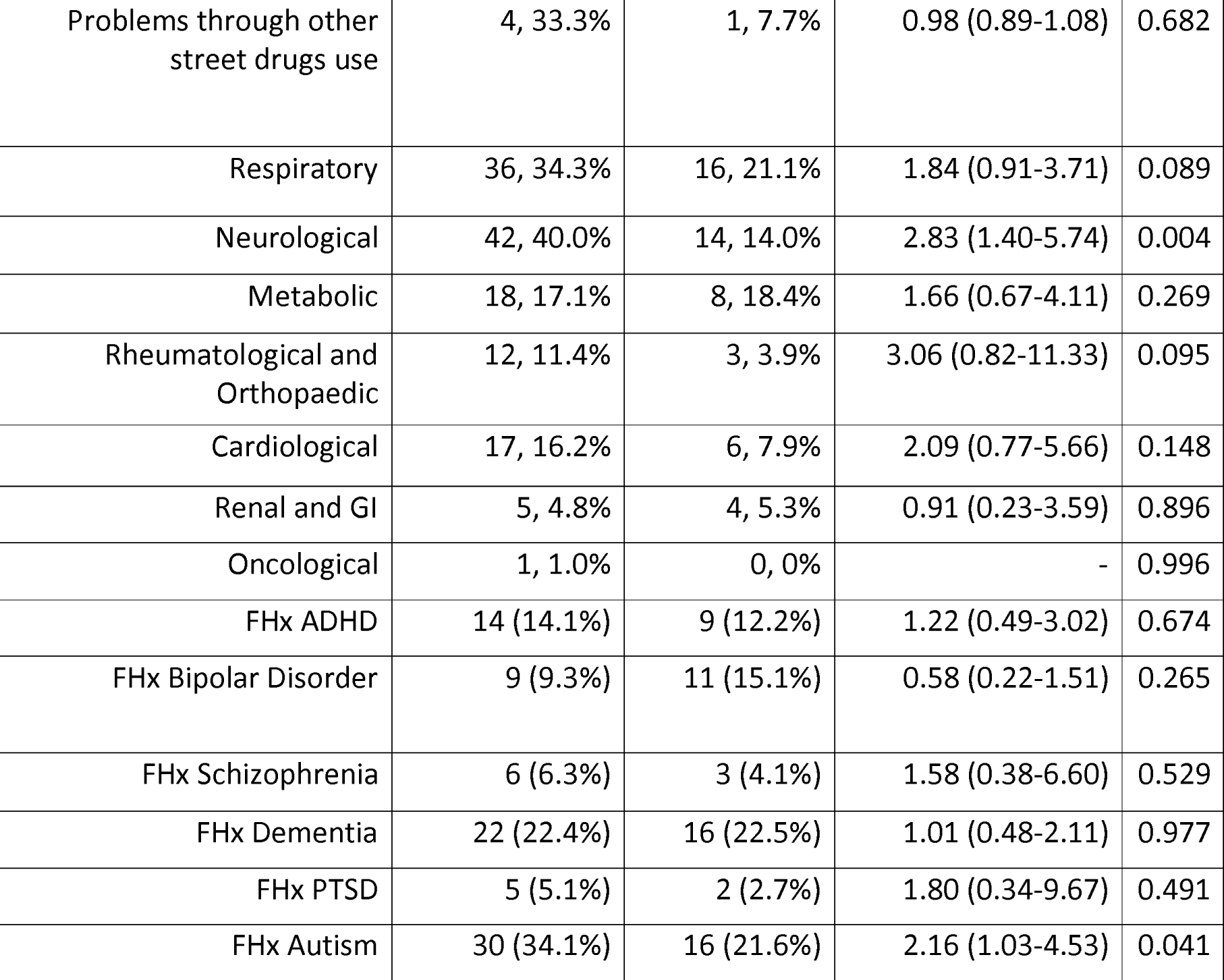
Physical health comorbidity, family history, social demographics and substance use in participants with ASD and matched controls Reported physical health comorbidity, family health history, social demographics and substance use in participants with ASD based on NCMH questionnaire response fields. The comparison group are matched healthy controls. For normally distributed samples, analyses were binomial logistic regression with age and sex as covariates. Where data were non-normally distributed, the non-parametric Mann-Whitney U test was used with Fisher’s exact test for expected cell counts <5. n - number of individuals; % - percentage of total who responded; FHx - reported family history.

|  | ASD participants (n,%) | Matched controls (n,%) | Odds Ratio (95% confidence interval) | p |
| --- | --- | --- | --- | --- |
| Currently Working | 31, 32.0% | 53, 72.6% | 0.174 (0.09-0.35) | <.001 |
| Currently not working due to sickness/ disablement | 48, 49.5% | 1, 0.01% | 69.305 (9.23-520.42) | <.001 |
| Biological Children | 36, 35.6% | 41, 54.0% | 0.526 (0.26-1.05) | 0.068 |
| Married or Cohabited | 49, 50.5% | 58, 76.3% | 0.29 (0.14-0.64) | 0.002 |
| Regular smoker | 34, 43.0% | 35, 46.1% | 1.017 (0.99-1.04) | 0.169 |
| Problems through alcohol use | 21, 36.2% | 5, 8.3% | 6.42 (2.20-18.72) | 0.001 |
| Regular cannabinoids use | 22, 27.9% | 16, 21.1% | 1.00 (0.97-1.03) | 0.903 |
| Problems through cannabinoid use | 9, 29.0% | 4, 16.0% | 1.02 (0.96-1.08) | 0.590 |
| Use of other street drugs | 10, 12.5% | 4, 5.3% | 0.99 (0.95-1.04) | 0.711 |
| Problems through other street drugs use | 4, 33.3% | 1, 7.7% | 0.98 (0.89-1.08) | 0.682 |
| Respiratory | 36, 34.3% | 16, 21.1% | 1.84 (0.91-3.71) | 0.089 |
| Neurological | 42, 40.0% | 14, 14.0% | 2.83 (1.40-5.74) | 0.004 |
| Metabolic | 18, 17.1% | 8, 18.4% | 1.66 (0.67-4.11) | 0.269 |
| Rheumatological and Orthopaedic | 12, 11.4% | 3, 3.9% | 3.06 (0.82-11.33) | 0.095 |
| Cardiological | 17, 16.2% | 6, 7.9% | 2.09 (0.77-5.66) | 0.148 |
| Renal and GI | 5, 4.8% | 4, 5.3% | 0.91 (0.23-3.59) | 0.896 |
| Oncological | 1, 1.0% | 0, 0% | - | 0.996 |
| FHx ADHD | 14 (14.1%) | 9 (12.2%) | 1.22 (0.49-3.02) | 0.674 |
| FHx Bipolar Disorder | 9 (9.3%) | 11 (15.1%) | 0.58 (0.22-1.51) | 0.265 |
| FHx Schizophrenia | 6 (6.3%) | 3 (4.1%) | 1.58 (0.38-6.60) | 0.529 |
| FHx Dementia | 22 (22.4%) | 16 (22.5%) | 1.01 (0.48-2.11) | 0.977 |
| FHx PTSD | 5 (5.1%) | 2 (2.7%) | 1.80 (0.34-9.67) | 0.491 |
| FHx Autism | 30 (34.1%) | 16 (21.6%) | 2.16 (1.03-4.53) | 0.041 |

### Physical Health Comorbidity

Adults with ASD were more likely to have neurological problems than controls (OR=2.83, 95% CI 1.40-5.74, p = 0.004). Posthoc analysis of the disorders within the neurological subgroup demonstrated this effect was predominantly due to an increased reported rate of migraine in individuals with ASD. Forty-one (42.7%) individuals with ASD reported lifetime history of migraine headaches compared to fifteen (20.5%) control participants (OR=2.60, 95% CI 1.24-5.44, p = 0.012). We subsequently explored whether the difference in these rates between ASD participants and controls could be explained by comorbid Epilepsy and Seizure Disorder, conditions reported co-occur at elevated rates paediatric ASD populations. This association between Migraine Headaches and Epilepsy and Seizure Disorders was confirmed (OR=11.38, 95% CI 1.31-99.09, p = 0.028).

### Family History of Psychiatric Disorder

Reported family history of autism was seen at a greater rate amongst the ASD participants (OR=2.16, 95% CI 1.03-4.53, p = 0.041), but no differences were found for family history of ADHD, Bipolar Disorder, Schizophrenia, Dementia or Post-traumatic Stress Disorder (PTSD).

## Genetic analysis

### CNV Analysis

The 53 neurodevelopmental CNVs examined for were present in 3.8% (n = 4) of ASD cases and 1.3% (n = 1) of controls. The CNVs in cases were 2q13 deletion (n = 2), 15q13.3 duplication (n = 1) and 16p13.11 duplication (n = 1). A single 2p16.3 deletion was found in a control.

### Analyses of Polygenic Risk Scores

Polygenic Risk Scores (PRS) derived from GWAS of ASD, ADHD, Alzheimer’s disease, Major Depressive Disorder, Bipolar Disorder and Schizophrenia were calculated for each individual. Only the PRS derived from the ASD GWAS showed differences between cases and controls (Figure 1); despite the modest sample size of this cohort, we calculate that approximately 12.86% (P < .00001) of variance can be explained from PRS derived from linkage independent markers showing association at P < 1e-3 in the ASD GWAS (SNPS in model = 433). No significant difference between cases and controls was seen for PRS for ADHD, Alzheimer’s disease, Major Depressive Disorder, Bipolar Disorder and Schizophrenia.

**Figure 1.**
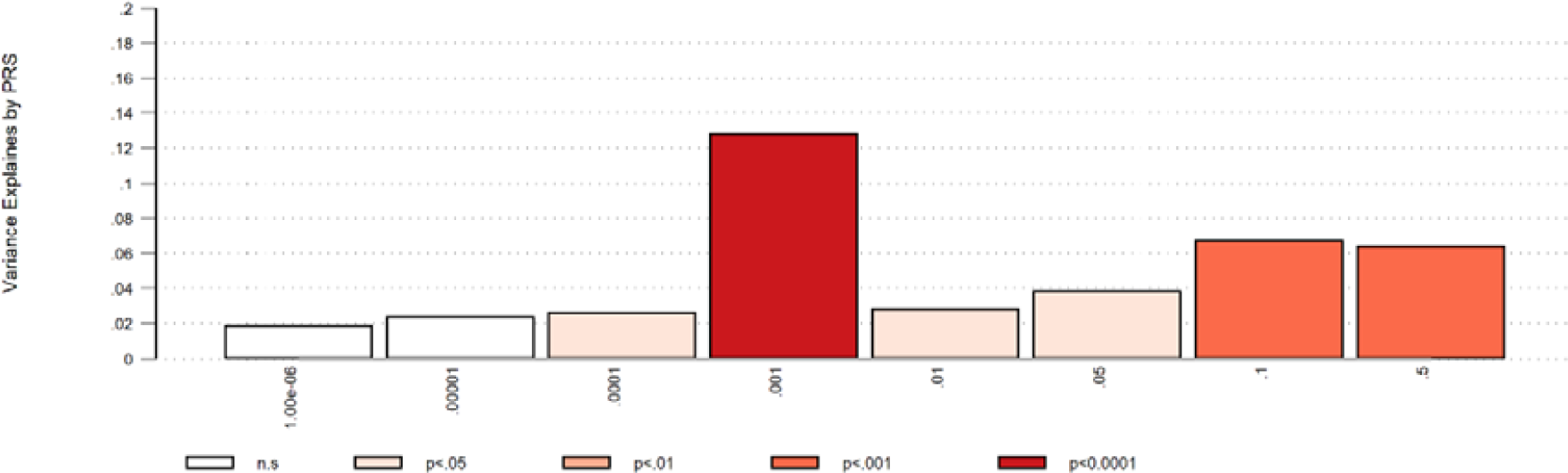
Percentage variance explained by PRS at analysed association levels for ASD. Percentage variance at eight association marker levels derived from linkage independent markers in the ASD GWAS. Significance of associations between SNPs and ASD range from 0.5 to 1 × 10^-6^. Probability of association to be found at each individual variance level is denoted by p value.

## Discussion

In this study we report on the phenotypic characteristics and genetic profiles of a sample of individuals with ASD diagnosed in adulthood without intellectual disability. We found high rates of psychiatric comorbidity, problem alcohol use and medication usage in individuals with ASD. These individuals also had higher rates of neurological comorbidity than controls and there was an association between ASD diagnosis and migraine. There was a significant association between PRS for ASD and a diagnosis of ASD but no significant increase in rate of rare neurodevelopmental CNVs in individuals with ASD.

### Comorbidity

89.5% of ASD participants reported a further lifetime psychiatric comorbidity, comparable to previously reported populations including those diagnosed as adults or with intellectual disability (69-80%)^7–10^. Depression was the most common lifetime comorbid diagnosis with a rate of 62.9%, comparable to previously reported rates (53-70%) in larger samples including those diagnosed as children or with intellectual disability ^7,11,30,31^. Lifetime anxiety diagnosis rates of around 50% were also similar to previously reported in those with ASD but with intellectual disability ^3,8–10,32^. The implication that a primary diagnosis of ASD is linked to high lifetime rates of anxiety and depression even in populations without ID is important for clinical consultations. Whether the aetiology of ASD predisposes for depression and anxiety, or life events and difficulties experienced by individuals because of their ASD precipitate depression and anxiety, or both, is a ‘chicken and egg’ conundrum worthy of further study. Rates of dyslexia (21.9%) and dyspraxia (18.1%) and ADHD (19.0%), while elevated compared to the general population, were lower than in previous studies of ASD which have mainly been in children ^5,7,33^. As our population were diagnosed as adults, fewer of the neurodevelopmental features which prompt assessment may be expected, and such a profile fits with genetic findings. Shared aetiology may explain the significantly higher rates of neurological conditions seen in the ASD participants. This is predominantly due to a strong prevalence of comorbid migraine. While a pathophysiological link has been suggested for this in the past, this is the first clear evidence of an association to our knowledge, and warrants further investigation ^34,35^.

Lifetime psychotropic medication usage was concordant with the diagnosis findings. Nearly 80% of the population had taken antidepressant medications during their lifetime, greater than the rates predicted in the literature ^9,32^. 26.3% reported taking antipsychotic medication when only 12.4% had a diagnosis of psychosis. This may be in part due to antipsychotic medication usage amongst four of the nine individuals reporting Bipolar Disorder, or for use off licence for symptomatic or behavioural management as is evidenced in adolescents and young adults ^9,36^.

### Social Demographics

The ∼3:1 male to female sex ratio seen in our population was consistent with reported gender differences ^2,5,33,37^. While alcohol and other substance use and smoking rates in our control and ASD participants were broadly similar, ASD was associated with higher rates of problematic alcohol use. The reasons were unclear, but could be usage to self-medicate for the aforementioned anxiety as suggested by other authors, or to facilitate social interactions ^5^.

Adults with ASD were found to be more than five times less likely to be employed individuals, with the vast majority reporting they were unemployed or unable to work due to illness. This suggests supportive employment is available for too few ^5^. We hypothesise that difficulties with social communication underlie the strikingly lower rate of marriage for those with ASD, although interestingly difference in rates of biological children are not significant. This points to the clear impact of ASD even when diagnosed in adulthood, and the contribution of comorbid psychiatric conditions to the life experience of an individual with ASD.

### Genetic Risk

Previous studies have demonstrated the strong heritability of ASD ^5^. As expected we demonstrated that adults with ASD were significantly more likely to have family members with ASD. There were no other significant associations with mental health disorder family history. This result may be of use for patients presenting to clinical practice wanting to know associated family risks but requires confirmation from a larger study. The ASD participants had a slightly higher number of neurodevelopmental disorder associated CNVs than controls, although this was not statistically significant in this sample. Although a statistically significant increase in burden of CNVs in this group might be confirmed in a larger population, rates were markedly lower than those reported in paediatric ASD populations, who may have a greater overall neurodevelopmental symptom profile ^29^. Polygenic risk score analysis demonstrated a significant contribution of polygenic load of ASD-associated common variants to risk in adult ASD participants ^15,24,38^. It is noticeable that a significantly increased polygenic risk burden was detected even in this relatively small sample size. Taken together these results suggest that adults presenting with ASD may have a lower burden of rare penetrant variants and a higher polygenic contribution of common risk alleles than childhood ASD populations, potentially reflecting less severe neurodevelopmental disability.

### Limitations

The sample used in this study were drawn from the NCMH database, a resource rich in phenotypic data and participants consented for anonymised genetic analysis ^16^. The benefit of the extensive range of data is offset by clinical diagnoses being initially self-reported, necessitating a review of the individuals’ notes to confirm specific diagnoses by specialist clinicians. All ASD diagnoses in this study were confirmed against ICD10 diagnostic criteria. This prevents analysis by symptom severity and lacks the clarity of diagnostic scoring systems or rating scales such as the ADI-R, however this reflects the pattern seen within mental health services where diagnoses are often clinical and diagnostic tool usage varies. By design our control population did not have psychiatric comorbidities, and this lack of control comorbidity data prevented comparative statistical analysis. All comorbid diagnoses were required to have been made by a doctor but were self-reported and therefore not standardised. As a descriptive study we benefited from the large quantity and extensive domains available from the NCMH dataset. Where possible results were elicited through subgroup analysis to reduce test volume. Associations here reported therefore require further testing with a more highly powered sample.

### Implications for services

In this study we provide comorbidity rates and social demographic information for adults presenting with ASD, with clinical utility for consultations in adult ASD diagnostic services. Our findings suggest that the vast majority of adults with ASD have psychiatric comorbidity and should be appropriately screened and managed. Additionally, clinicians should be aware of associated social demographic features including the high rates of alcohol problem use and being out of work. Signposting towards and integration with third sector organisations and services supporting adults with ASD is vital, and our results may inform these services towards the possible difficulties some diagnosed with ASD in adulthood may face. The significantly increased polygenic risk burden as seen in our sample is a difficult concept to convey in a clinical environment. It is likely that for some individuals genetic testing would provide an element of assurance and diagnostic clarity, and our results may assist in genetic counselling. For many the benefit of an ambiguous answer is questionable. This is an area where future work with larger population sizes promises better results. Our findings suggest that the polygenic risk burden is present in clinical samples, and ongoing advances may allow us to explain this fully along with the associations and implications.

**There are no conflicts of interest to declare.**

## Authorship Credit

Underwood, Jack F G – contributed at all stages of the study and manuscript.

Kendall, Kimberley M – contributed to all stages of the study and manuscript.

Berrett, Jennifer – contributed to all stages of the study and manuscript.

Anney, Richard – contributed to analysis and interpretation of polygenic risk score data, drafting of the manuscript and approval.

Van den Bree, Marianne B.M – contributed to analysis of statistical data, drafting of the manuscript and approval.

Hall, Jeremy – contributed to all stages of the study and manuscript.

## Acknowledgements

The authors would like to thank Professor Ian Jones and Dr Catrin Lewis for their assistance and enthusiasm, along with the rest of the team at NCMH, the Bioinformatics team at Cardiff University and all the staff at MRC CNGG. The NCMH is a collaboration between Cardiff, Swansea and Bangor Universities and is funded by Welsh Government through Health and Care Research Wales. We thank the NCMH study participants for their invaluable contribution to this project.

